# A synthetic approach reveals a highly sensitive maize auxin response circuit

**DOI:** 10.1101/844100

**Authors:** Román Ramos Báez, Yuli Buckley, Han Yu, Zongliang Chen, Andrea Gallavotti, Jennifer L. Nemhauser, Britney L. Moss

## Abstract

Auxin plays a key role across all land plants in growth and developmental processes. Although auxin signaling function has diverged and expanded, differences in the molecular functions of signaling components have largely been characterized in *Arabidopsis thaliana*. Here, we used the Auxin Response Circuit recapitulated in *Saccharomyces cerevisiae* (ARC^Sc)^ system to functionally annotate maize auxin signaling components, focusing on genes expressed during development of ear and tassel inflorescences. All 16 maize Auxin (Aux)/Indole-3-Acetic Acid (IAA) repressor proteins are degraded in response to auxin, with rates that depended on both receptor and repressor identity. When fused to the maize TOPLESS (TPL) homolog RAMOSA1 ENHANCER LOCUS2 (REL2), maize Aux/IAAs were able to repress AUXIN RESPONSE FACTOR (ARF) transcriptional activity. A complete auxin response circuit comprised of all maize components, including ZmAFB2/3 b1 maize AUXIN SIGNALING F-BOX (AFB) receptor, was found to be fully functional. The ZmAFB2/3 b1 auxin receptor was found to be more sensitive to hormone than AtAFB2 and allowed for rapid circuit activation upon auxin addition. These results validate the conserved role of predicted auxin response genes in maize, as well as provide evidence that a synthetic approach can facilitate broader comparative studies across the wide range of species with sequenced genomes.

A synthetic maize auxin response circuit is recapitulated in Saccharomyces cerevisiae, revealing a highly sensitive auxin signaling network with functional homology to the Arabidopsis circuit.

## INTRODUCTION

Auxin is an ancient molecule, and its role as a phytohormone dates back to the earliest diverging land plants (Mutte et al., 2018). The expansion of the gene families that encode the auxin response pathway parallels increasing complexity in plant form. For example, the moss *Physcomitrella patens* has three members of the auxin (Aux)/Indole-3-Acetic-Acid (IAA) gene family (Prigge et al., 2010), while the eudicot *Arabidopsis thaliana* (*Arabidopsis*) has 29 and the monocot *Zea mays* (maize) has 34 (Ludwig et al., 2013; Luo et al., 2018; Matthes et al., 2019). The retention of large gene families involved in auxin response following genome duplications, in combination with the central role of auxin in plant development, has led to the hypothesis that functional diversification in auxin response circuits underpins structural and functional innovations during land plant evolution (Mutte et al., 2018). Connecting natural variation to functional divergence remains a major challenge. This problem becomes even more complicated if selection is operating on the amplitude or dynamics of a network rather than the function of any one component, a premise that itself has been quite difficult to test.

The auxin response is among the best understood signaling pathways in plants, and thus is an excellent model to tackle questions about functional divergence in gene families within a single species and in network functions across multiple species. Auxin response consists of three modules: (i) activation of auxin responsive genes by AUXIN RESPONSE FACTOR (ARF) transcription factors; (ii) repression of ARFs in the absence of auxin by recruitment of TPL/TOPLESS-RELATED (TPR) co-repressors via Aux/IAA proteins; and (iii) degradation of Aux/IAA proteins via their auxin-induced association with TRANSPORT INHIBITOR RESPONSE1 (TIR1)/AUXIN SIGNALING F-BOX (AFB) receptors (Leyser, 2018). While there is some evidence for distinct combinations of these proteins acting in different cell types (Bargmann et al., 2013; Rademacher et al., 2012; Vernoux et al., 2011), co-expression of components, feedback regulation and interactions with other signaling pathways (Piya et al., 2014) have made it virtually impossible to conclusively support or refute this model.

The wealth of genetic, genomic and biochemical tools in *Arabidopsis* have made it possible to rapidly build a strong foundational understanding of auxin response. However, to both explore the extent of shared auxin signaling properties across plants and to fully interrogate the connection between natural variation in protein sequences and functional innovations in plant development, the auxin response must be examined in more species. Understanding differences between eudicots and monocots is of particular interest, as the molecular mechanism that explains the differential impacts of widely-used auxinic herbicides remains a mystery, where eudicot weeds are killed while grasses are often unaffected (McSteen, 2010). In recent years, functional studies in maize have made in-roads in delineating auxin response components. For example, the maize Aux/IAA protein RUM1, is known to be functionally important in the development of the root in the embryo, as well as root branching in seedlings (von Behrens et al., 2011). BIF1 and BIF4 are Aux/IAAs known to have important functions in the development of maize inflorescences (Galli et al., 2015). Mutations in REL2, a functional homolog of TPL in *Arabidopsis*, have pleiotropic auxin developmental effects in maize, rice, and *Arabidopsis* and can rescue TPL mutants in *Arabidopsis* (Gallavotti et al., 2010; Liu et al., 2019; Yoshida et al., 2012). Maize ARFs show similar preferences for auxin response elements (AuxREs) as *Arabidopsis* ARFs (Galli et al., 2018).

Auxin response circuits built in yeast cells (ARC^Sc^) make it possible to analyze individual and combinatorial functions of Aux/IAA, TIR1/AFB, ARF and TPL/TPR proteins (Havens et al., 2012; Pierre-Jerome et al., 2014). This heterologous system has several advantages, including precise control over the amount and duration of auxin input, highly quantitative fluorescence outputs, and the ability to study auxin signaling modules with defined connectivity and in the absence of other plant signaling pathways. Studies using *Arabidopsis* components in ARC^Sc^ have shown that Aux/IAAs exhibit a range of tunable auxin-induced degradation rates and that this variation in Aux/IAA degradation is central to controlling auxin transcriptional response dynamics and the rate of developmental events *in planta* (Guseman et al., 2015; Havens et al., 2012; Moss et al., 2015; Pierre-Jerome et al., 2014). The yeast system has also enabled functional characterization of genetic natural variation in auxin receptors (Wright et al., 2017) and functional annotation of putative maize auxin repressor proteins and mutant variants (Galli et al., 2015).

Here, we have used the ARC^Sc^ system to functionally annotate maize auxin signaling components, focusing on genes that are expressed during inflorescence development. All maize Aux/IAAs (ZmIAAs) tested degraded in response to auxin, with rates that depended on both receptor and repressor identity. ZmIAAs were able to repress ARF-mediated transcription when fused to a truncated form of the maize TPL homolog REL2. The maize auxin receptor ZmAFB2/3 b1 was found to be highly sensitive to auxin, allowing for more rapid degradation of Aux/IAAs than what was observed with the fastest acting Arabidopsis receptor AtAFB2. Finally, a complete auxin response circuit comprised of all maize components was found to be fully functional, allowing highly sensitive activation of a transcriptional reporter following addition of auxin. These results validate the conserved role of predicted auxin response genes in maize, as well as provide evidence that a synthetic approach can facilitate broader comparative studies that incorporate the wide range of species with sequenced genomes.

## RESULTS

### ZmIAAs are Functional in Auxin Degradation and Repression Modules

The maize B73 genome contains 34 Aux/IAA repressor genes, 16 of which were selected for further analysis here because they are expressed in developing maize inflorescences (Bolduc et al., 2012; Davidson et al., 2011; Eveland et al., 2014) and are representative of the phylogenetic diversity of the Aux/IAA family (Eveland et al., 2014; Matthes et al., 2019). Out of 13 ZmAux/IAAs that were tested by *in situ* hybridizations, six gave detectable and specific expression patterns in immature tassel inflorescence meristems and spikelet meristems (Fig. 1A-B). In general, most of these *ZmAux/IAA* genes revealed highly-restricted expression patterns marking emerging primordia in inflorescence meristems (*ZmIAA2, ZmIAA5, ZmIAA9, ZmIAA14, ZmIAA28)* as well as vasculature (i.e. *ZmIAA5* and *ZmIAA28*), and all showed strong expression in spikelet meristems (Fig. 1B). These expression patterns strongly resembled those previously reported for *BIF1* and *BIF4* (Galli et al., 2015) and suggest a high degree of functional redundancy in this family of transcriptional regulators. However, it is not known whether auxin-dependent degradation dynamics or repression strengths may reveal differences in molecular function among individual *Aux/IAAs*. To assess the auxin sensitivity of each ZmIAA, we fused these inflorescence-expressed ZmIAAs to YFP and co-expressed them with an Arabidopsis auxin receptor in Saccharomyces cerevisiae (yeast) (Fig. 1C). All ZmIAA proteins were expressed in yeast cells and degraded in response to auxin treatment (Fig. 1D). The ARC^Sc^ auxin degradation module also allowed us to measure functional differences in auxin-induced degradation rates by measuring changes in fluorescence following auxin exposure (Havens et al., 2012). We selected three ZmIAAs that were expressed at similar levels and showed different auxin sensitivities (ZmIAA8, ZmIAA12, and BIF1) and measured auxin-induced degradation dynamics (Fig. 1E). As seen previously for AtIAAs (Havens et al., 2012), ZmIAA degradation rate varied between different ZmIAAs (Fig. 1E), and was generally more rapid in cells expressing AtAFB2 compared with those expressing AtTIR1 (Fig. 1F).

**Fig. 1.**
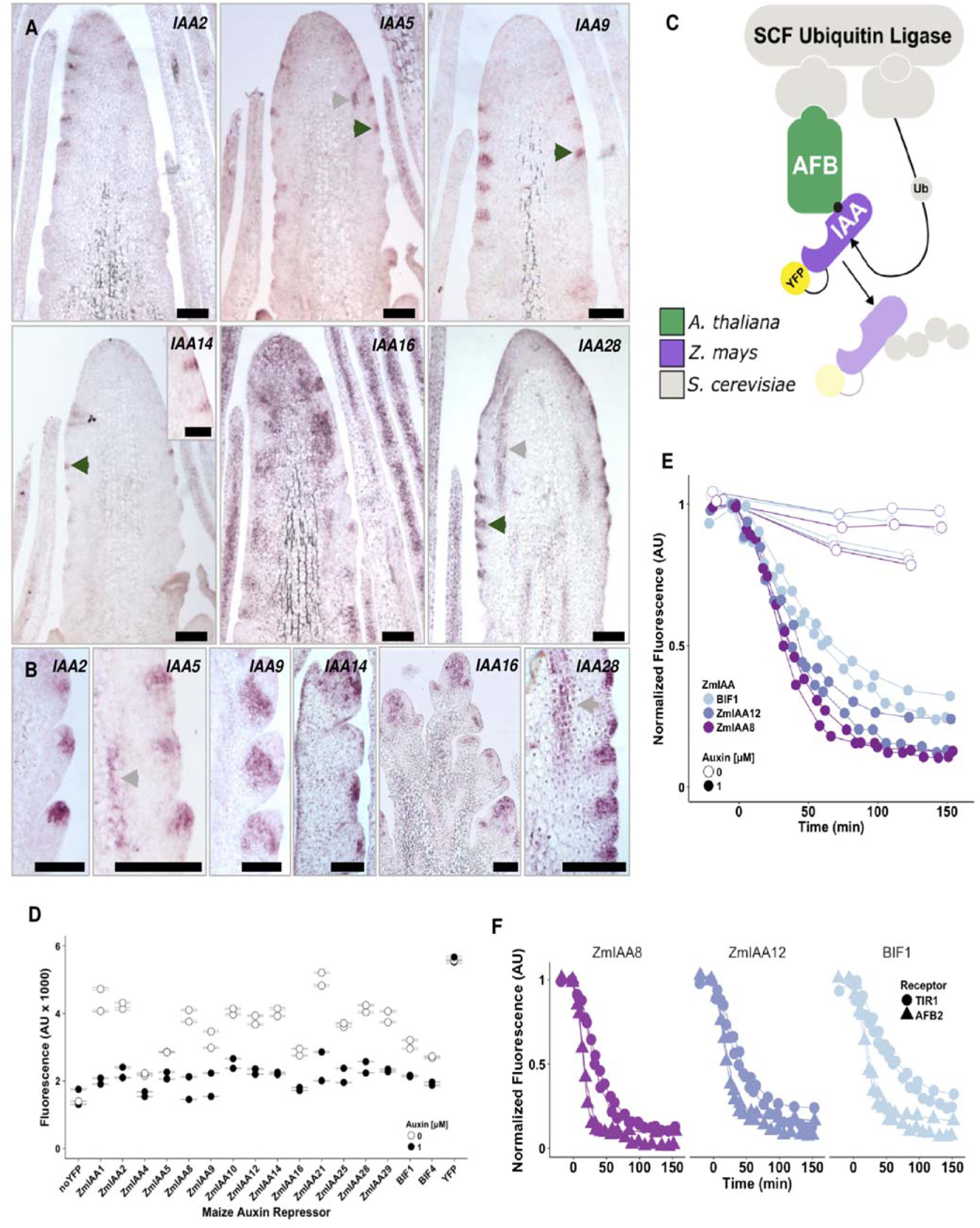
Auxin repressors expressed in maize inflorescence exhibited variable auxin-induced degradation dynamics. **(A,B)** Expression of maize *Aux/IAA* genes by mRNA *in situ* hybridizations in immature tassels. Expression patterns in inflorescence meristems (**A**) and spikelet meristems (**B**). Grey arrows, vasculature; black arrows, axillary meristems. Scale bars, 100 μm in (**A**) and (**B**) and 50 μm in inset picture. **(C)** The auxin degradation module in yeast consists of ZmIAAs (purple) tagged with YFP and co-expressed with an Arabidopsis auxin receptor (green). The F-box domain of the receptor facilitates complex formation with the yeast SCF Ubiquitin Ligase machinery (gray). When yeast are exposed to auxin the hormone acts as molecular glue that brings the co-receptor complex together and results in ubiquitination and degradation of the YFP-tagged ZmIAA, resulting in a decrease in fluorescence over time. **(D)** The 16 ZmIAAs were degraded in response to auxin. Fluorescence measurements on flow cytometer were obtained 2 hours post-auxin exposure. Data from two replicates shown, error bars represent SEM. **(E,F)** ZmIAAs degrade at different rates that are dependent upon both repressor **(E)** and receptor identities **(F)**. Yeast strains expressing YFP-tagged ZmIAAs and either *Arabidopsis* TIR1 or AFB2 auxin receptor were exposed to 1 μM auxin or mock treatment (95% ethanol) at 0 min and fluorescence measurements taken on flow cytometer. Data from two replicates shown.

In addition to degradation in response to auxin, the other major function of Aux/IAAs is repression of ARF-mediated transcriptional regulation, a process that is facilitated by TPL/TPR co-repressors (Causier et al., 2012). The maize genome has four TPL-like co-repressors, the REL2/REL2-like (RELK) family, of which REL2 has been shown to have pleiotropic phenotypes associated with meristem maintenance and initiation in maize (Liu et al., 2019). In the Arabidopsis ARC^Sc^ (AtARC^Sc^), an N-terminal fusion of TPLN100 was found to be functional if directly fused to IAAs, facilitating transcriptional repression of ARFs (Pierre-Jerome et al., 2014). We confirmed that REL2 can also confer repression of ZmIAAs by fusing ZmIAA8 to either TPLN100 or REL2N91, a fragment of REL2 that is structurally analogous to TPL100 (Fig. 2A & S3C). Based on new structural information, we designed REL2N91 to include only the first five helices that encompass the LisH and CTLH domains (Martin-Arevalillo et al., 2017). The ARC^Sc^ strains used for repression assays contained Arabidopsis AFB2 and ARF19, which was shown to be the strongest and fastest activating ARF in AtARC^Sc^ (Pierre-Jerome et al., 2014). Each co-repressor conferred a similar degree of repression, and in the presence of auxin that repression was relieved, and transcription was activated to similar degrees (Fig. 2B). Both the degree of repression and auxin-induced activation dynamics varied greatly across ZmIAAs, and repression level did not necessarily predict activation level (Fig. 2C, S1B). Two ZmIAAs were unable to repress the auxin response (Fig. S1C), possibly due to poor expression or inability to interact with ARF19.

**Fig. 2.**
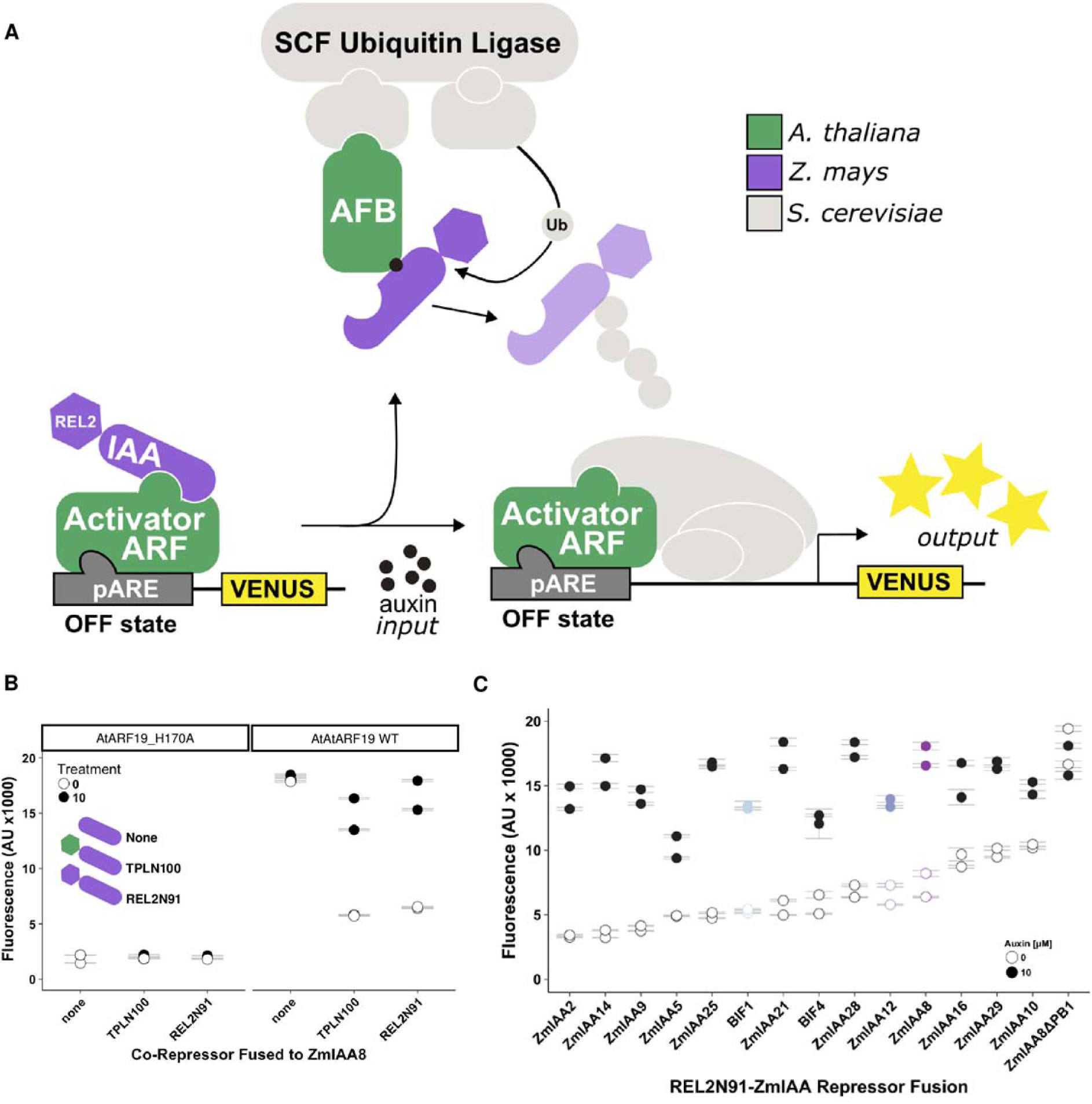
The TPL homolog REL2 enabled *Zm*IAAs to repress ARFs. **(A)** The auxin repression module in yeast consisted of *Zm*IAA repressors fused to N-terminal fragments of either the *Arabidopsis* TPL or maize REL2 co-repressors; these were co-expressed with an auxin receptor (*Arabidopsis* AFB2) and activator transcription factor (*Arabidopsis* ARF19). Auxin-induced de-repression of the ARF19 results in activation of the auxin response element-containing promoter (pARE) and expression of VENUS fluorescent protein. **(B)** The N-terminal 91 amino acids of maize REL2 assist *Zm*IAA8 in conferring transcriptional repression on ARF19, and this repression is relieved upon addition of auxin. The REL2N91 fragment was directly compared to the analogous *Arabidopsis* TPLN100. The *At*ARF19_H170A mutant is unable to bind DNA and so auxin response stays off. Strains labeled “none” contain a *Zm*IAA8 that has not been fused to a co-repressor. **(C)** *Zm*IAAs fused to REL2N91 exhibited different patterns of auxin responsive gene activation, independent of their basal repression strength and degradation rate. Two *Zm*IAAs were unable to repress the auxin response (Fig. S1C). All fluorescent measurements were made 5 hours post-treatment. Data from two replicates shown, error bars represent SEM.

### Maize Auxin Receptor ZmAFB2/3 b1 is Highly Sensitive to Auxin

In the maize genome, there are eight members of the TIR1/AFB auxin receptor family (Matthes et al., 2019), four of which appear to be related to the Arabidopsis AFB2/3 clade. Two of these AFB2/3-like maize auxin receptors were highly expressed in inflorescence meristems and so were tested for activity within an ARC^Sc^ degradation module (Bolduc et al., 2012; Eveland et al., 2014). The ZmAFB2/3 b1 receptor, was the only one able to induce degradation of ZmIAAs in the presence of auxin (Fig. 3A). This receptor also happens to be the most highly expressed in developing inflorescences (Eveland et al., 2014). Alignments and pairwise comparisons between receptors suggested they all share about 60% sequence similarity across all domains (Fig. S2A, B-D). Auxin-induced degradation carried out by ZmAFB2/3 b1 was even faster than AtAFB2 (Fig. 3A). We next assessed expression of the auxin receptors in the yeast strains and found that all were expressed and accumulated to similar levels (Fig. 3B, S2E-F), suggesting that differences in expression level are not likely to explain differences in degradation module activity. Furthermore, the amount of AtAFB2 receptor accumulating in yeast (within a two-fold range) was previously shown to have no effect on auxin-induced repressor degradation rate (Havens et al., 2012).

**Fig. 3.**
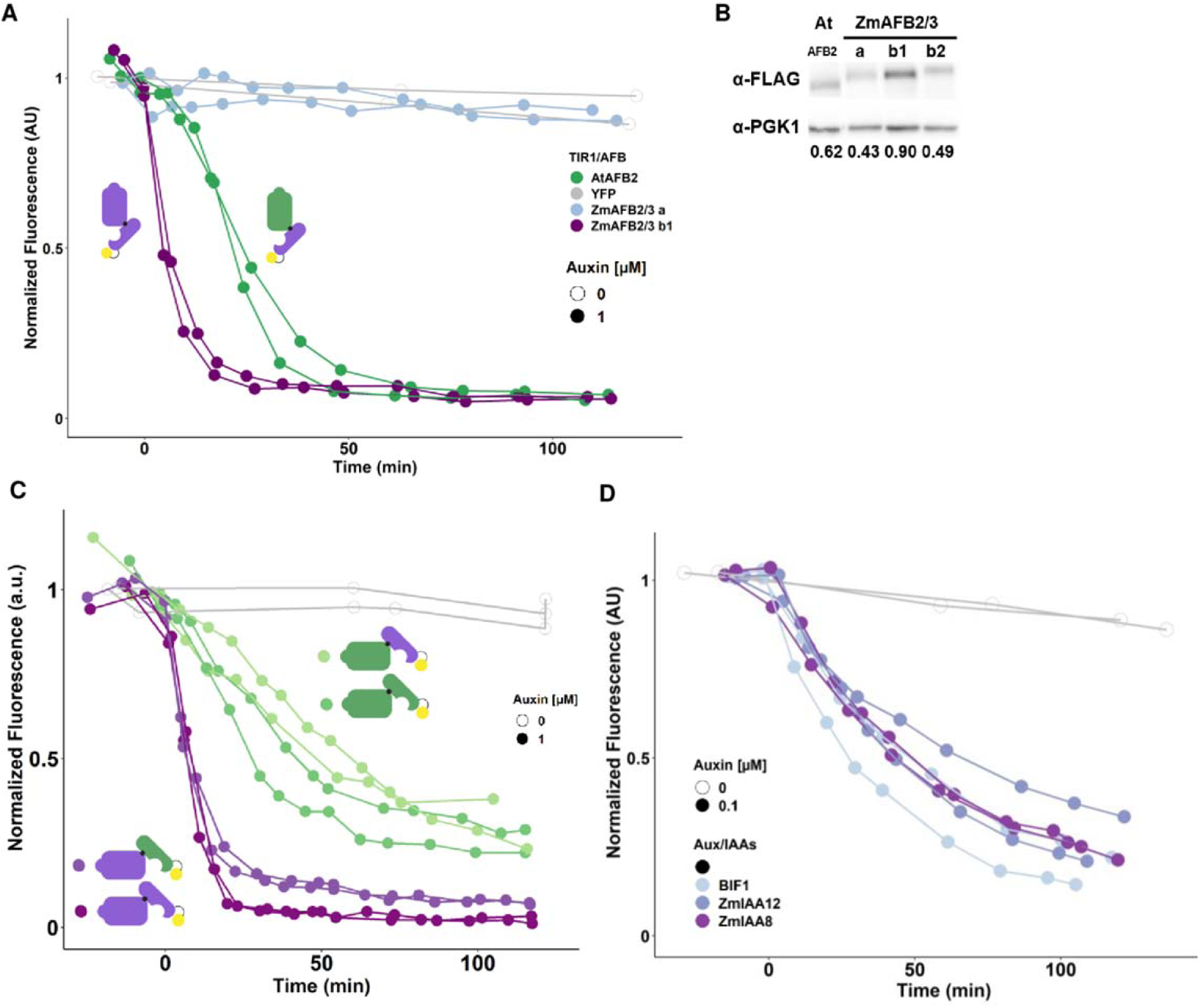
The maize auxin receptor ZmAFB2/3 b1 exhibited a higher basal activity compared to Arabidopsis ortholog. **(A)** ZmAFB2/3b1 was sensitive to auxin but had diminished dynamic sensitivity compared to AtAFB2. Yeast co-expressing an Arabidopsis or maize auxin receptor and YFP-ZmIAA10 were treated with 1 μM auxin or mock (“0”) and fluorescence was measured on a flow cytometer. **(B)** Auxin receptor expression level in yeast does not necessarily correlate with functionality. Yeast lysates were probed with anti-Flag (for receptors) or anti-PGK1 (loading control) antibodies. Fold-expression values shown below bands were calculated by using ImageJ to quantify the intensity of each band and dividing the intensity of the receptor band by the intensity of the PGK1 band. **(C)** ZmAFB2/3b1 always exhibited faster IAA degradation in response to auxin, whether paired with an Arabidopsis or a maize Aux/IAA. All combinations of ZmAFB2/3b1 or AtAFB2 paired with AtIAA14 or ZmIAA14 were tested for auxin degradation module behavior. **(D)** When paired with ZmAFB2/3b1, three maize Aux/IAAs (BIF1, ZmIAA8, and ZmIAA12) exhibited different orders of degradation speed than when paired with AtAFB2 (see Fig. 1).

The auxin sensitivity of ZmAFB2/3 b1 and AtAFB2 were next tested with Aux/IAAs from each species to determine which component was primarily responsible for driving differences in speed of the degradation module. The maize receptor was faster than AtAFB2 when paired with either a maize or an Arabidopsis Aux/IAA (Fig. 3C, Fig. S4B). We also noticed that pre-auxin Aux/IAA accumulation levels were lower in strains expressing ZmAFB2/3 b1 compared to AtAFB2 (Fig. S3A). We hypothesized that the maize receptor might be interacting with the repressor independently of auxin. Treatment with auxinole, a compound shown to block interaction of rice TIR1 (OsTIR1) and auxin repressors (Yesbolatova et al., 2019), resulted in greater accumulation of Aux/IAAs, particularly in yeast strains expressing ZmAFB2/3 b1 (Fig. S3A). This suggested that at least some of the increased auxin sensitivity of ZmAFB2/3 b1 may be due to auxin-independent degradation. Auxin-sensitivity analyses showed that ZmAFB2/3 b1 is responsive to auxin over two orders of magnitude (0.01 – 1.0 μM), (Fig. S3B). We next assessed whether ZmAFB2/3 b1 degradation modules were functional with each of the 16 ZmIAAs. All ZmIAAs were degraded by ZmAFB2/3 b1 (Fig 3D, S3C). Furthermore, some ZmIAAs exhibited different patterns of auxin-induced degradation dynamics than observed when co-expressed with AtAFB2: for example, BIF1 paired with ZmAFB2/3b1 degraded as fast or faster than ZmIAA8 or ZmIAA12 when paired with AtAFB2 (Fig. 1E, 3D; notice also the 10-fold difference in auxin sensitivities). This is reminiscent of the different patterns of ZmIAA degradation rates observed for Arabidopsis AFB2 and TIR1 auxin receptors (Havens et al., 2012). Thus, maize auxin degradation modules are highly sensitive to hormone, and degradation dynamics are dependent upon both receptor and repressor identity.

### A Fully-Maize Auxin Response Circuit is Tuned to Respond to Low Auxin Concentrations

Having determined that maize receptors, repressors, and co-repressors are functional in varying combinations, the final step was to express them with a maize auxin transcription factor to reconstitute an all-maize auxin full circuit (ZmARC^Sc^, Fig. 4A). ZmARF27 is orthologous to AtARF19, the ARF used in the AtARC^Sc^, and is expressed strongly in immature tassel (Eveland et al., 2014; Matthes et al., 2019). ZmARC^Sc^ yeast strains containing various ZmIAAs were able to repress ZmARF27 transcriptional activity to varying degrees (Fig. 4B), conferring as strong or stronger repression on ZmARF27 than on AtARF19 (Fig. S4A). The ZmARC^Sc^ strains were all de-repressed by the addition of a high concentration of auxin (Fig. 4B, S4A). Given that ZmAFB2/3 b1 is more sensitive to auxin than AtAFB2 in auxin degradation modules (Fig. 3), we hypothesized that this would also result in ZmARC^Sc^ transcriptional responses activating at lower doses of auxin. For each ZmARC^Sc^ strain tested, the transcriptional response was activated at lower auxin levels compared to strains in which the ZmAFB2/3b1 was replaced with AtAFB2 (Fig. 4C, S4B). Thus, the maize auxin response circuit is exquisitely-sensitive to auxin hormone levels and response dynamics can be tuned by altering ZmIAA identity.

**Fig. 4.**
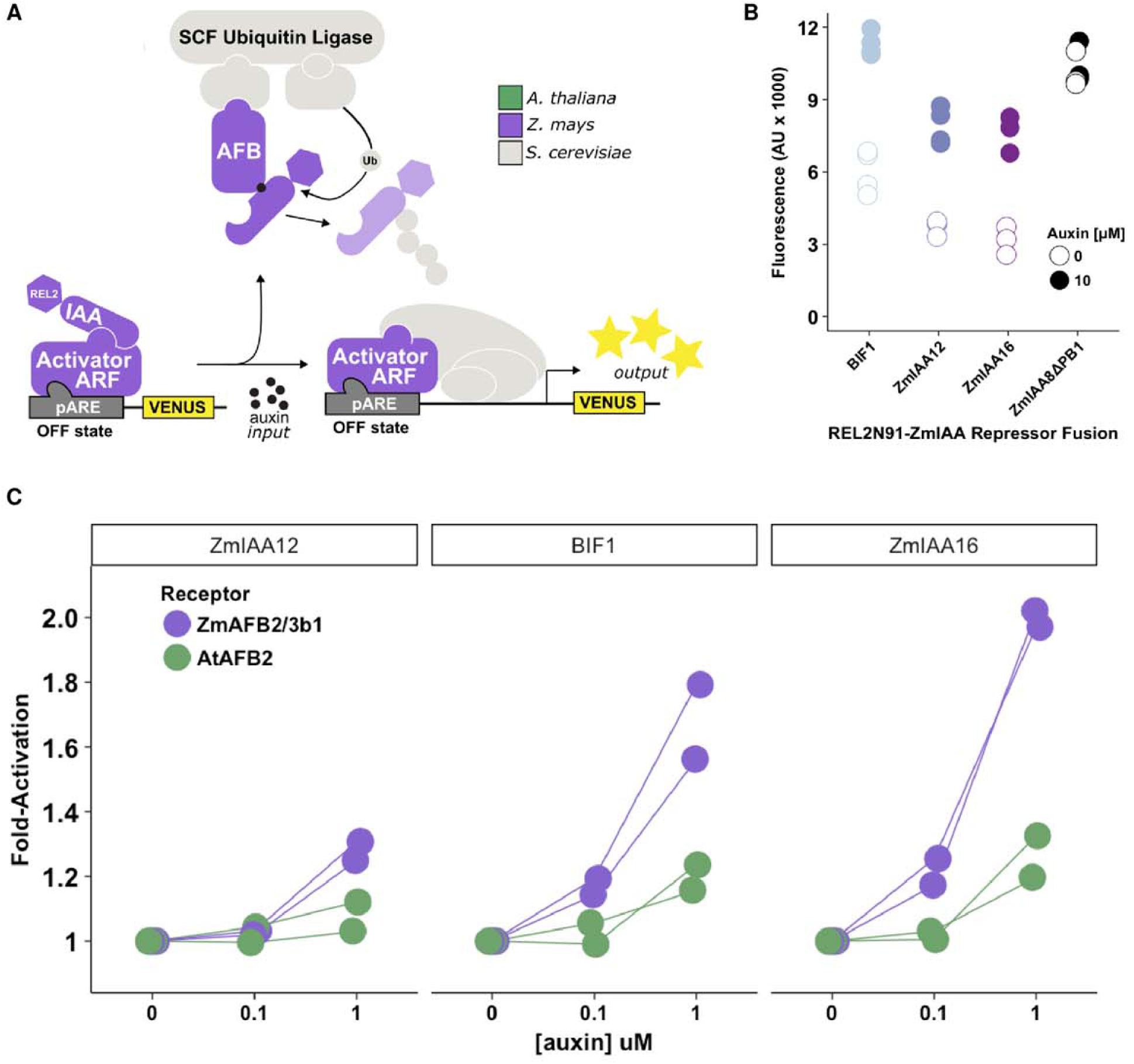
Fully-maize auxin response circuit is functional and highly sensitive to auxin. **(A)** A fully-maize auxin response circuit in yeast (ZmARC^Sc^) was assembled as shown with ZmARF27. **(B)** The fully-maize auxin response circuit is functional and responds to auxin. Data represents 3-4 replicates of fluorescence measurements taken 5 hours following hormone treatment. The ZmIAA8 strain with a deleted PB1 domain again represents an always “on” state. **(C)** ZmAFB2/3b1 auxin receptor confers high auxin-sensitivity to the maize auxin response circuit. Strains shown in (B) and their AtAFB2 cognates were treated with 0, 0.1, or 1 μM auxin for 5 hours before fluorescence measurements. Data plotted as fold-untreated.

## DISCUSSION

The synthetic auxin response circuit used in this study is impressively modular, with the ability to mix-and-match components from distantly-related plant species. Maize ZmIAAs expressed in developing inflorescence tissues degraded in response to auxin, with variable degradation rates that depended on both ZmIAA and receptor identity (Fig. 1). Furthermore, these ZmIAAs were able to repress transcription when paired with the maize co-repressor REL2 (Fig. 2). Fully-maize auxin degradation modules were found to be more sensitive to auxin than orthologous Arabidopsis degradation modules (Fig. 3). Finally, an all-maize auxin response circuit featuring *Zm*ARF27 was fully functional and more sensitive to auxin than circuits containing an *Arabidopsis* receptor (Fig. 4). This work represents a broadly useful strategy for employing synthetic biology approaches to functionally annotate and characterize genetic diversity within conserved signaling networks.

Decreasing costs of genome sequencing have led to a dramatic increase in genetic natural variation information available. However, functional annotation of this new genetic information is lagging behind. The ARC^Sc^ platform allows for rapid functional validation and characterization of large gene families, as seen previously for auxin signaling modules within *Arabidopsis* (Havens et al., 2012; Moss et al., 2015; Pierre-Jerome et al., 2017, 2014; Wright et al., 2017) and now for maize (Fig. 1 & 2). Of particular note, the yeast system enabled multi-dimensional functional annotation (degradation and repression behaviors) of sixteen different Aux/IAAs expressed in developing maize inflorescences. We observed variable degradation and repression behaviors across the ZmIAAs studied, providing evidence of biochemical differences that introduce another layer of complexity on top of known phylogenetic diversity (Ludwig et al., 2013) and tissue-specific expression patterns (Matthes et al., 2019). In agreement with previous studies (Ludwig et al., 2013; von Behrens et al., 2011; Zhang et al., 2016), ZmIAA2, ZmIAA10/RUM1, ZmIAA20/BIF4, and ZmIAA29/RUL1 were shown to be functional within auxin degradation modules (Fig. 1 & 3), and capable of repressing transcription (Fig. 2). Further, by showing that ZmIAAs can repress the auxin circuit with the assistance of REL2, we provide an additional piece of evidence to support existing genetic and biochemical data demonstrating that REL2 is a functional homolog of TPL (Liu et al., 2019). Thus, our functional annotation and characterization of maize auxin signaling modules is in agreement with the few existing genetic and biochemical studies on maize Aux/IAAs and REL2, and extends functional characterization to many previously uninvestigated maize auxin signaling components for which there are no known mutants. The recent recapitulation of the *Arabidopsis* ABA hormone signaling pathway in yeast underscores the feasibility of performing similar functional annotation efforts for putative orthologous proteins across many plant signaling pathways (Ruschhaupt et al., 2019).

The ARC^Sc^ system is a valuable platform for continued dissecting of the differences in auxin response between monocots and dicots. For example, ARC^Sc^ should be useful in elucidating the mechanisms of auxinic herbicide tolerance and resistance from the perspectives of chemical biology, evolution, and genetic engineering. In monocots, responses to most auxinic herbicides are much weaker than in eudicots, which die when these herbicides are applied to them (McSteen, 2010). Auxin biosynthesis and transport pathways are largely conserved suggesting that differences in auxin signaling and reception in monocots might be what primarily contributes to these different responses. Because maize and *Arabidopsis* both have similar number of genes in auxin signaling gene families (TIR1/AFBs, Aux/IAAs, ARFs, etc.) we believe differences between auxin-dependent development in these species are likely driven more by differences in protein function, not differences in pathway protein stoichiometries. Our results bring up exciting new questions about the role of auxin in maize. ZmAFB2/3 b1 is more sensitive to lower auxin dosages than AtAFB2 and can lead to Aux/IAA turnover even in the absence of auxin (Fig. 3). Analysis of relative auxin levels during tassel and ear development, perhaps using the DII-VENUS reporter (Mir et al., 2017), could provide clues to the impact of this increased sensitivity on feedback within the auxin system and/or on the dynamics of downstream responses.

Understanding how hormone signaling protein functions have changed (or remained similar) throughout evolution can help connect protein function to growth and developmental processes and inform the design of synthetic proteins. ARC^Sc^ has revealed the functional ramifications of the evolution of auxin signaling components within *Arabidopsis*, allowing for the functional comparison of gene variants across accessions (Wright et al., 2017). By allowing for quantitative comparisons between *Arabidopsis* and maize auxin signaling pathways, ARC^Sc^ is a promising tool to study the evolution of molecular function more broadly across distantly related plant species, in this case diverging 250MYA. It also provides the ability to directly and quickly study molecular functions in multiple dimensions to help elucidate the evolution of wide-spread developmental phenomena such as inflorescence development. From a synthetic biology perspective, platforms like ARC^Sc^ can enable rapid design, characterization, and comparison of highly divergent proteins for use in engineering synthetic signaling machinery within plants or other biological systems.

The types of experiments and analyses described here that utilize a synthetic hormone signaling system in yeast are readily carried out by novice scientists in research laboratories and in course settings. Many of the experiments described in this paper were piloted or executed by 45 undergraduate students at a Primarily Undergraduate Institution, Whitman College. Most of these students were participants in several semesters of course-based undergraduate research experiences (CUREs). This type of undergraduate-focused synthetic biology research experience offers an exciting way to acquaint young scientists with plant synthetic biology, and could be readily adapted to focus on standardization and characterization of parts for use in synthetic biology, an area of continuing concern (Decoene et al., 2018).

## METHODS

### *in situ* Hybridizations in maize inflorescences

For mRNA *in situ* hybridizations, 0.2–0.4 cm tassel primordia from the B73 inbred line were dissected and fixed in cold paraformaldehyde acetic acid (PFA) solution as described previously (Galli et al., 2015). Hybridizations were carried out at 59°C. Antisense *in situ* probes for all thirteen *AUX/IAA* genes (*IAA2*, *IAA4*, *IAA5*, *IAA8*, *IAA9*, *IAA10*, *IAA12*, *IAA14*, *IAA16, IAA21, IAA25*, *IAA28* and *IAA29*) were synthesized by *in vitro* transcription (T7 RNA polymerase, Promega) of the entire or partial coding sequences cloned in pENTR vector or pGEM T-easy (Promega) and digested with respective endonuclease enzymes. The vectors and primers used for probe design are listed in Table ST01.

### Yeast Methods

Yeast (*Saccharomyces cerevisiae*) were grown in either yeast peptone dextrose (YPD) or Synthetic Complete (SC) media made according to standard recipes and supplemented with 80 mg/mL adenine. Yeast transformation of linearized plasmids was performed using a standard lithium acetate protocol (Gietz and Woods, 2002). Matings were performed by co-inoculation of MATa and MATɑ strains at low density in YPD media with shaking overnight at 30°C. All yeast transformations and matings were streaked onto SC plates lacking appropriate auxotrophic compounds for selection, followed by isostreaking onto YPAD plates prior to glycerol stocking. Full yeast strain list is in Table ST02.

### Plasmid Construction

Plasmid and primers design was performed in Benchling. Maize *Aux/IAA* and *ARF* sequences were obtained from Grassius Database or synthesized (IDT). Maize REL2 fragment was cloned from plasmids generated in (Liu et al., 2019). The maize *TIR1/AFB* gene sequences were obtained from Paula McSteen’s lab. These sequences were obtained from IDT with codon optimization for *S. cerevisiae*, and then cloned into pCR-BLUNT plasmid using the Zero Blunt® TOPO® PCR Cloning Kit (James et al., 2000). The *ZmIAA* genes were inserted into pGP4GY plasmids (Havens et al., 2012) via Gateway cloning (Life Technologies), or by Gibson cloning (Gibson et al., 2009). *ZmARF27* was cloned into the pGP8G vector by Gibson cloning. Auxin receptors (*TIR1* and *AFB2s*) were cloned into pGP8GF plasmid containing an upstream 3-Flag, 6-HIS tag. Each PCR amplification was performed using Q5® High-Fidelity DNA Polymerase (New England Biolabs), and the products were purified using the EZNA Cycle Pure® Kit or the NEB Monarch® PCR & DNA Cleanup Kit and confirmed by sequencing (MCLab or Genewiz).

### Auxin Degradation Module Assays

Auxin degradation module assays were carried out as previously described (Pierre-Jerome et al., 2017). In brief, yeast colonies from YPAD plates were used to inoculate SC media. Cell density (in events/μL) was estimated by flow cytometry and cultures were diluted such that cells were in log-phase growth 16 hr later and for the duration of the assay. All growth was at 30°C in an incubated shaking at 375 rpm. Fluorescence measurements were taken on the flow cytometer prior to the addition of auxin to establish baseline. Auxin (indicated concentrations, indole-3-acetic acid) or mock (95% [v/v] ethanol) treatments were added, and cultures were returned to incubator. Fluorescence measurements were acquired at 10 min intervals for the first hour after auxin addition and every 15 min thereafter until the fluorescence level in most strains had plateaued (approximately 2 hrs). Controls strains were measured every hour for the duration of the experiment. For auxinole (50 μM in ethanol) treated strains, fluorescence measurement was taken 4 hrs following treatment.

### Repression/Activation Module Assays

Auxin repression/activation module assays were carried out as previously described (Pierre-Jerome et al., 2017). Briefly, yeast cells were grown overnight in 2 mL SC media at 30°C in an incubated shaker. In the morning, the cells were diluted 1:200 into fresh SC and returned to the incubator. After two hours, auxin (indicated concentrations) or mock treatment (95% [v/v] ethanol) were added and strains continued to incubate at 30°C for 5 hrs. Fluorescence measurements were taken on flow cytometer.

### Flow Cytometry

Fluorescence measurements were taken with a BD Accuri C6 flow cytometer and CSampler plate adapter using an excitation wavelength of 488 and an emission detection filter at 533 nm. A total of 10,000 - 20,000 events above a 400,000 FSC-H threshold (to exclude debris) were collected for each sample and data exported as FCS 3.0 files for processing using the flowCore R software package and custom R scripts (Havens et al., 2012; Pierre-Jerome et al., 2017). Data from at least two independent replicates were combined and plotted in R.

### Western Blotting

Yeast were grown overnight in 5mL liquid YPD at 30°C with shaking at 225 rpm. The next morning cultures were diluted to an OD of 0.2 into fresh YPD media and grown until an OD of ~1.0. Cells were then pelleted and resuspended in 1% SDS, 8M Urea, 10mM MOPS, pH 6.8, 10mM EDTA, 0.01% bromophenol blue (SUMEB) buffer with 1mM phenylmethylsulfonyl fluoride (PMSF). Cell pellets were lysed by glass bead disruption in lysis buffer containing protease inhibitors. Aliquots of 7µL of yeast proteins in buffer were loaded onto 10% SDS-PAGE gel (Bio-Rad). Anti-FLAG (Cell Signaling Technology) was used to probe for TIR1/AFBs-3XFLAG6XHIS, and anti-PGK1 (Invitrogen) was used as a loading control.

## ACKNOWLEDGEMENTS

Thank you to members of the Nemhauser and Imaizumi labs for helpful discussion and guidance on experimental design and execution. Thank you to Paula McSteen and Norman Best for advice for providing maize receptor sequences and for advice on the manuscript. Thank you especially to the students in Britney Moss’s Synthetic Cell Biology classes, Tristan Cates, Kristina Jackson, Kathleen Daly-Jensen, Hannah Klaeser, and Alex Koriath for initial experimental work. This work was supported by the National Science Foundation IOS-1546873 (to J.N., A.G., B.M.) and IOS-1456950 (to A.G.), the M.J. Murdock Charitable Trust (to B.M.), and Whitman College (to B.M.). Román Ramos Báez was supported by the Howard Hughes Medical Institute Gilliam Fellowship GT11355.

## SUPPLEMENTAL MATERIALS

Supplemental Table 1 ST01 and

Supplemental Table 2 ST02 “SSpreadsheets RAMOS BAEZ 2019.xlsx”

Supplemental Figure 1 S1 and

Supplemental Figure 2 S2 and

Supplemental Figure 3 S3 and

Supplemental Figure 4 S4 “SFigures RAMOS BAEZ 2019.pdf”

**Supplemental Figure 1. The N-terminal REL2 portion fused to *Zm*IAAs represses transcription. (A)** TPLN100 and REL2N91 were aligned using MegaX software. They show high sequence similarity. Only positions 40 and 43-48 are different. **(B)** ZmIAAs fused to REL2N91 were inserted into the repression module. These show a range of activation and repression levels, measured with flow cytometry for accumulation of and auxin-responsive UbiVENUS. The purple box highlights the two ZmIAAs that did not repress transcription. **(C)** Timecourse of transcriptional activation in strains with REL2N91-ZmIAA fusions. Auxin or mock treatment (95% Ethanol) was added at t = 0 min and fluorescence measurements were made on a flow cytometer.

**Supplemental Figure 2. The maize auxin AFB2/3 receptors proteins are orthologous to those in other species and expressed at similar levels in yeast. (A)** Alignment of the amino acid sequences of functional Arabidopsis, maize, and rice TIR1/AFB receptors using MegaX Clustal Omega, and visualized using CLC Workbench. **(B)** Full-length amino acid sequence pairwise comparisons of Arabidopsis and maize functional TIR1/AFBs show high sequence similarity. **(B)**. The similarities are conserved across both the F-box domains **(C)**, and the LRR domains **(D)**. Two reps each of auxin receptor Western blots showing **(E)** PGK1 protein loading controls and **(F)** anti-FLAG blots showing different AFB2/3 protein quantities in yeast lines. Positive control was OsAFB2-3F6H-expressing yeast, which had previously been shown to be functional and expressed in our degradation module. Negative control was yeast without an integrated transgene.

**Supplemental Figure 3. ZmAFB2/3 b1 degraded ZmIAAs more readily in the absence of auxin, and was more sensitive to auxin. (A)** Fully-maize degradation modules expressing either ZmIAA21 or 28 were checked for fluorescence using cytometry. A range of auxin concentrations were added at time zero. **(B)** Auxinole was added to cultures with degradation modules with Arabidopsis and maize components. These cultures were then diluted, and their fluorescence was measured 6hrs post-auxinole addition. Auxinole addition resulted in higher IAA concentrations in all lines, but especially in ZmAFB2/3 b1-containing lines. **(C)** All ZmIAAs tagged to YFP were measured for degradation after auxin addition at time zero. Fluorescence was measure for two hours. Two reps.

**Supplemental Figure 4. Auxin full circuit behavior depended on the identity of the ARF, TIR1/AFB, and IAA (A)** Circuits expressing different ARFs showed different levels of repression, as well as different levels of activation post auxin-addition. **(B)** Circuits expressing AtAFB2 or ZmAFB2/3 b1 had different sensitivities to auxin, and different basal levels of repression.

